# BirdAVES in the wild: individual recognition as a step toward zebra finch communication networks

**DOI:** 10.1101/2025.11.20.689459

**Authors:** Aneesh Chauhan, Hugo Loning, Evelien ter Avest, Bhavya Botta

## Abstract

Understanding who communicates with whom, when, and how is central to the ecology of group-living animals, yet individual-level acoustic identification of animals in their natural environment remains challenging. Zebra finches are a model species whose vocal behaviour has been predominantly studied indoors; here we address the outdoor setting and investigate bioacoustic deep learning for individual identification at scale as a key step to build communication networks from field recordings. We fine-tune BirdAVES for recognizing 173 zebra finch individuals from short (1-3 s) clips using a concise training recipe: two-phase training, weighted sampling and class-weighted cross-entropy for long-tailed counts, and a supervised contrastive term to pull same-individual embeddings together. On a real-world dataset (2,915 clips, 173 individuals), the selected model achieved macro-F1 = 0.733 (val) / 0.726 (test) and steep retrieval gains (Top-5 = 0.868, Top-10 = 0.893 on test set). This enables conversion of hours of audio into “who-sang-when” timelines. We deliberately report top-k performance because it quantifies review effort and supports human-in-the-loop workflows by shrinking the number of clips an expert must audit. While a train–val/test gap reflects short windows and class imbalance, the embeddings are discriminative and immediately useful. Key next steps are to address the imbalance in our data and scaling towards a significantly larger set of individuals, and to translate individual recognition into communication or social networks.

## 1 Introduction

The zebra finch (Taeniopygia castanotis) is one of the most-studied songbird in the lab [5], but it remains challenging to ecologically contextualise the resulting wealth of lab-based knowledge. In the wild, zebra finches are non-territorial songbirds, living in dynamic fission–fusion groups [19]. Their vocalisations are very short-distance signals, with an average detection range of less than 14 meters [11]. Thus, vocalisations are not efficient in locating and attracting conspecifics in the open, arid habitat zebra finches live in. Instead, zebra finches use temporally stable gathering locations, so-called ‘social hotspots’ to interact with conspecifics [12]. It has been hypothesized that their individually distinct vocalisations [4] might facilitate the individual recognition necessary to establish and maintain social connections [13]. Proximity-based social networks can be obtained using passive [3] and active [16] tags, but these tags cannot measure how and how much individuals are actually interacting. Using acoustic data to recover so-called communication networks at the level of individuals would enable a significantly better understanding about social organization, information flow, and coordination of this model species. However, the key prerequisite - robust, scalable identification of individuals from acoustic data - remains challenging.

Large-scale bioacoustic deep learning is evolving along two complementary tracks [9]: (i) *supervised* species-level classifiers (e.g., BirdNET [8]; Perch[17]) trained on large-scale labeled data, and (ii) *label-efficient* representation learners, like *self-supervised* foundation models (e.g., BirdAVES [6], Animal2Vec [15]) that learn without annotations and often transfer better with limited labels. Supervised models excel where dense species labels exist (common birds), but their coverage degrades for rare/endangered taxa and non-avian groups with limited data. In these low-label scenarios, self-supervised pretraining provides stronger initializations and more generalizable embeddings. Motivated by this, we adopt pretrained BirdAVES model as the choice for zebra finch individual identification.

Our approach builds on progress in bioacoustic modelling, in particular a combination of BirdAves and supervised contrastive learning [10] and is complementary to prototype-based contrastive learning [14]. Our dataset comprises 173 individuals, each with multiple recordings. The class distribution is highly imbalanced, ranging from a single recording to 177 recordings for an individual. Furthermore, besides noise, field audio from non-territorial species, like the zebra finch, introduces specific hurdles: short-range signals with overlapping callers yield sparse, local views; data are imbalanced across individuals and contexts. We leverage BirdAVES and adapt it into a multi-class classifier. We train the classifier using a concise recipe: two-phase training, weighted sampling and class-weighted cross-entropy to address the data imbalance, and a supervised contrastive loss [10] to pull same-individual embeddings together while preserving class separation.

## 2 Methodology

### 2.1 Dataset

#### Data collection and annotation

Primary fieldwork took place at Fowlers Gap (NSW, Australia) from 4 Sep. to 28 Oct. 2023. We placed SM3 song meters (Wildlife Acoustics) in hotspot trees, recording 16 kHz/16-bit mono WAV continuously from sunrise to sunset. We analyzed four hotspots: five days per site for three hotspots ( 65 h per tree) and three days for a fourth, highly active hotspot (198 h total). Audio files were screened in Audacity (v3.4.2) via spectrograms (1024-sample Hann, 0–8 kHz). Annotation proceeded in two stages: (1) detect song bouts (vs. calls); (2) label song bouts to individuals (only males sing and songs are individually distinctive). Zebra finch songs were identified by introductory notes followed by one or more motifs. An expert then assigned song bouts to individuals using reference exemplars; a second expert reviewer was consulted for uncertain cases. Disagreements remained unlabeled. We expanded this dataset corpus with prior (2018) handheld recordings from the same site, adding measured song motifs (which may be shorter than full bouts). For preprocessing, segments >6 s were trimmed to the first and last 3 s; segments *≤* 3*s* were left unchanged.

#### Dataset composition

Our dataset comprises 2,915 audio segments (clips) from 173 individuals. Segments are short and largely fixed-length (median 3 s, min 0.7 s, max 3 s). Per-individual counts are highly imbalanced (median 8 clips; max 177), with 14 individuals represented by a single clip. We split the dataset by clip (identities may appear in multiple partitions; some do not appear in val/test due to low counts) and ensure no identical recording is reused across splits. Composition by clips: train 69.9% (2,038), val 17.6% (513), test 12.5% (364). Identity coverage: 173/173 in train, 116/173 (67.1%) in val, 100/173 (57.8%) in test.

### 2.2 Modelling approach

We define individual identification as a 173-way multi-class classification problem from short zebra finch audio clips.

#### Architecture

A pretrained BirdAVES [6] (Wav2Vec2 [2] / HuBERT-based [7]) encoder produces frame-level features; we mean-pool them into a 1024-D embedding and add a linear head to produce 173 logits. This converts BirdAVES into a 173-class classifier that outputs raw logits.

#### Input data preprocessing

Waveforms are mono, resampled to 16 kHz, and windowed to 3.0 s by zero-padding. Because HuBERT-style encoder (as used in the BirdAVES) is sensitive to padding, validation uses batch size 1 (no cross-sample padding).

#### Handling class imbalance

We combine (i) *oversampling* via a WeightedRandomSampler, PyTorch with sampling *s*(*i*)*∝* 1*/n*_*c*_, where *n*_*c*_ is the number of training clips for individual *c*; and (ii) *class-weighted* cross-entropy with

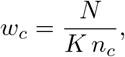

where *N* is the number of training clips and *K* the number of individuals in the training data.

#### Objective and optimization

We optimise over two loss functions. (i) *ℒ*_CE_, class-weighted cross-entropy, and (ii) *ℒ*_SupCon_, supervised contrastive loss [10] on *l*2 *−normalised* embeddings. Our implementation scales the averaged log-probability by temperature *τ* = 0.07, rescaling *ℒ*_SupCon_ and is absorbed by the contrastive weighting *λ* below. When a batch contains only one sample (e.g., validation with batch size 1), the contrastive term is disabled. We minimise the combined loss

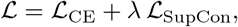

using Adam optimizer with separate learning rates for the encoder and the classifier head and use gradient accumulation (ACCUM_STEPS = 8).

Training is divided into two pahses. Phase 1 minimises *ℒ*_CE_ with the encoder frozen; Phase 2 minimises *ℒ* with the encoder unfrozen. Here, *λ* balances classification and contrastive representation learning, while *τ* controls the sharpness of contrastive softmax (smaller *τ* enforces tighter positive clusters and larger angular margins).

#### Hyperparameter search

We tune for batch size [4, 20], head/encoder learning rates [10^*−*7^, 10^*−*4^] (log scale), epochs_1_ [5, 30], epochs_2_ [30, 200], and *λ* [0.1, 0.9] using Optuna [1](median pruning). Runs are tracked with MLflow [18]; models and configurations are saved per trial.

#### Training infrastructure

All experiments were conducted on an NVIDIA GeForce RTX 3090 GPU (24 GB) with PyTorch 2.3.1 (CUDA 11.8).

**Evaluation protocol and metric**

Data are split by clip (individuals will appear across splits but no clip is duplicated). Given our highly imbalanced data, we report by macro-F1, which averages per-class F1 and therefore reflects performance on both rare and frequent individuals. We use macro-F1 as the objective to be maximized for the Optuna runs.

## 3 Results

From the Optuna runs, we select the model with the best macro-F1 on the validation set. The best model had the following training parameters: batch size = 14, epochs = 21 (Phase 1), 62 (Phase 2), learning rates: head lr_*head*_ = 1.02*×* 10^*−*7^, encoder lr_*enc*_ = 9.14*×*10^*−*5^ (Phase 2), contrastive weight *λ* = 0.513.

Table 1 reports the results corresponding to this model. We observe a clear train–val/test gap (macro-F1≈ 0.99 vs. 0.73). This indicates *some* level of overfitting, but is also consistent with (i) severe per-individual data imbalance and (ii) distributional differences across splits (e.g., many individuals with few clips in val/test). Macro-F1, which weighs each identity equally, amplifies this effect. However, the *top-k* curves rise steeply (e.g., test Top-5 = 0.868, Top-10 = 0.893), suggesting the representation is discriminative but the Top-1 decision is brittle for low-support identities.

**Table 1:** Overall performance of the selected checkpoint. Macro-F1 is computed over classes present in each split. We report Top-k accuracies to quantify *how far down the ranked list* the correct identity appears - useful for shortlist-based verification (human-in-the-loop).

| Split | Macro-F1 | Top-1 | Top-3 | Top-5 | Top-10 | Top-20 | Top-30 | Top-50 |
| --- | --- | --- | --- | --- | --- | --- | --- | --- |
| Train | 0.9919 | 0.9818 | 0.9975 | 0.9985 | 0.9990 | 0.9990 | 0.9995 | 0.9995 |
| Val | 0.7332 | 0.7485 | 0.8577 | 0.8850 | 0.9181 | 0.9376 | 0.9454 | 0.9513 |
| Test | 0.7255 | 0.7445 | 0.8434 | 0.8681 | 0.8929 | 0.9258 | 0.9341 | 0.9451 |

Top-k accuracies measure how far down the model’s ranking the correct individual appears. This is useful when a small shortlist enables fast human verification. Top-k counts a prediction correct if the true class is among the top k logits. In our results the curves rise quickly, so the correct identity is usually within a short shortlist.

Per-class Top-*k* accuracy statistics on the test set are reported in Table 2, with the corresponding distributions shown in Figure 1. The median Top-1 accuracy is 1.0, indicating that for at least half of the classes the model consistently predicts the correct label on the first attempt. This trend persists across all *k*, reflecting a substantial subset of “easier” classes that are reliably recognized. In contrast, the relatively large standard deviation at Top-1 (≈ 0.36) highlights considerable variability across classes, with some classes rarely or never identified correctly. As *k* increases, the deviation decreases (reaching 0.21 at Top-50), suggesting that harder classes benefit from more permissive evaluation while easier classes remain saturated at perfect accuracy.

**Table 2:** Per-class Top-*k* summary (test set).

| Top-$k$ | Mean | Median | Std | Min |
| --- | --- | --- | --- | --- |
| Top-1 | 0.739 | 1.0 | 0.358 | 0.0 |
| Top-3 | 0.826 | 1.0 | 0.306 | 0.0 |
| Top-5 | 0.844 | 1.0 | 0.298 | 0.0 |
| Top-10 | 0.856 | 1.0 | 0.296 | 0.0 |
| Top-20 | 0.905 | 1.0 | 0.248 | 0.0 |
| Top-30 | 0.917 | 1.0 | 0.230 | 0.0 |
| Top-50 | 0.934 | 1.0 | 0.208 | 0.0 |

**Figure 1:**
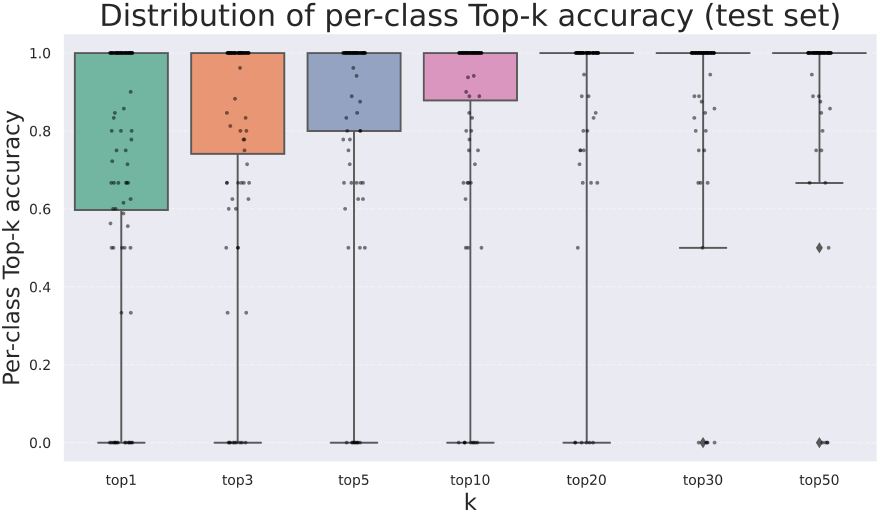
Distribution of per-class Top-*k* accuracies (test set).

Overall, the results demonstrate that while strict Top-1 recognition remains challenging under severe class imbalance, the model achieves strong shortlist retrieval performance, making it practical for downstream ecological analyses, from supporting individual-level inference to communication or social network reconstruction in wild zebra finch populations.

## 4 Conclusions

Bioacoustic deep learning can play an important role in studying zebra finch communication networks at scale, and in the wild. Zebra-finches have predominantly (historically) been studied indoors under controlled conditions, here we tackle outdoor field audio from wild birds in their natural habitat. Individual recogntion in the wild will allow us to quantify zebra finch song bouts across multiple contexts - e.g., hotspots vs. breeding sub-colonies - and to compare the resulting communication networks. While songs are short-range signals and thus provide local snapshots, those snapshots are precisely what is needed to connect vocal behaviour to proximity-based social structure.

Here we show that a concise fine-tuning recipe: BirdAVES encoder with a 173-way head, two-phase training, weighted sampling + class-weighted cross-entropy, and a supervised contrastive term - can deliver actionable individual recogntion performance. Our best model achieves macro-F1 = 0.733 (val) / 0.726 (test) with steep retrieval gains (Top-5 = 0.868, Top-10 = 0.893 on the test set). This can enable long recordings to be converted into *who-sang-when* timelines. Reporting top-*k* is also deliberate: it supports *human-in-the-loop* workflows by shrinking the number of clips an expert must review per decision. This moves individual-level inference beyond the lab and toward constructing communication networks in natural settings.

A measurable train–val/test gap (∼0.99 vs. ∼0.73 macro-F1) indicates some overfitting which is largely amplified by long-tailed per-individual counts (highly imbalanced data per individual, ranging anywhere from 1 clip to 177, with a median of 8 per individual). A practical approach is to create more balanced datasets, which is a work in progress for us as we record and annotate new data. However, the most important open problem is identifying new (unseen) individuals in the wild. Solving this will make individual recognition robust at population scale and can help create automated communication and social networks as a standard ecological tool. This problem is not limited to our research, but broadly applicable to bioacoustic deep learning research.

## Acknowledgments

This research was primarily funded by the Ministry of Agriculture, Fisheries, Food Security and Nature (LVVN) in the Netherlands via the Knowledge Base programme *Technological Innovations for Ecosystem Monitoring*, grant nr. KB-52-000-005.

## References

[1] Takuya Akiba, Shotaro Sano, Toshihiko Yanase, Takeru Ohta, and Masanori Koyama. Optuna: A next-generation hyperparameter optimization framework. In Proceedings of the 25th ACM SIGKDD international conference on knowledge discovery & data mining, pages 2623–2631, 2019.

[2] Alexei Baevski, Yuhao Zhou, Abdelrahman Mohamed, and Michael Auli. wav2vec 2.0: A framework for self-supervised learning of speech representations. Advances in neural information processing systems, 33:12449–12460, 2020.

[3] Hanja B Brandl, Simon C Griffith, Damien R Farine, and Wiebke Schuett. Wild zebra finches that nest synchronously have long-term stable social ties. Journal of Animal Ecology, 90(1): 76–86, 2021.

[4] Julie E Elie and Frédéric E Theunissen. Zebra finches identify individuals using vocal signatures unique to each call type. Nature communications, 9(1):4026, 2018.

[5] Simon C Griffith and Katherine L Buchanan. The zebra finch: the ultimate australian supermodel. Emu, 110(3):v–xii, 2010.

[6] Masato Hagiwara. Aves: Animal vocalization encoder based on self-supervision. In ICASSP 2023-2023 IEEE International Conference on Acoustics, Speech and Signal Processing (ICASSP), pages 1–5. IEEE, 2023.

[7] Wei-Ning Hsu, Benjamin Bolte, Yao-Hung Hubert Tsai, Kushal Lakhotia, Ruslan Salakhutdinov, and Abdelrahman Mohamed. Hubert: Self-supervised speech representation learning by masked prediction of hidden units. IEEE/ACM transactions on audio, speech, and language processing, 29:3451–3460, 2021.

[8] Stefan Kahl, Connor M Wood, Maximilian Eibl, and Holger Klinck. Birdnet: A deep learning solution for avian diversity monitoring. Ecological Informatics, 61:101236, 2021.

[9] Vincent S Kather, Burooj Ghani, and Dan Stowell. Clustering and novel class recognition: evaluating bioacoustic deep learning feature extractors. arXiv preprint 2504.06710, 2025.

[10] Prannay Khosla, Piotr Teterwak, Chen Wang, Aaron Sarna, Yonglong Tian, Phillip Isola, Aaron Maschinot, Ce Liu, and Dilip Krishnan. Supervised contrastive learning. Advances in neural information processing systems, 33:18661–18673, 2020.

[11] Hugo Loning, Simon C Griffith, and Marc Naguib. Zebra finch song is a very short-range signal in the wild: evidence from an integrated approach. Behavioral Ecology, 33(1):37–46, 2022.

[12] Hugo Loning, Rita Fragueira, Marc Naguib, and Simon C Griffith. Hanging out in the outback: the use of social hotspots by wild zebra finches. Journal of Avian Biology, 2023(11-12):e03140, 2023.

[13] Hugo Loning, Simon C Griffith, and Marc Naguib. The ecology of zebra finch song and its implications for vocal communication in multi-level societies. Philosophical Transactions B, 379(1905):20230191, 2024.

[14] Ilyass Moummad, Romain Serizel, Emmanouil Benetos, and Nicolas Farrugia. Domain-invariant representation learning of bird sounds. arXiv preprint 2409.08589, 2024.

[15] Julian C Schäfer-Zimmermann, Vlad Demartsev, Baptiste Averly, Kiran Dhanjal-Adams, Mathieu Duteil, Gabriella Gall, Marius Faiß, Lily Johnson-Ulrich, Dan Stowell, Marta B Manser, et al. animal2vec and meerkat: A self-supervised transformer for rare-event raw audio input and a large-scale reference dataset for bioacoustics. arXiv preprint 2406.01253, 2024.

[16] Chris Tyson, Hugo Loning, Simon C Griffith, and Marc Naguib. Constant companions: wild zebra finch pairs display extreme spatial cohesion. Biology Letters, 20(11):20240519, 2024.

[17] Bart van Merriënboer, Vincent Dumoulin, Jenny Hamer, Lauren Harrell, Andrea Burns, and Tom Denton. Perch 2.0: The bittern lesson for bioacoustics. arXiv preprint 2508.04665, 2025.

[18] Matei A. Zaharia, Andrew Chen, Aaron Davidson, Ali Ghodsi, Sue Ann Hong, Andy Konwinski, Siddharth Murching, Tomas Nykodym, Paul Ogilvie, Mani Parkhe, Fen Xie, and Corey Zumar. Accelerating the Machine Learning Lifecycle with MLflow. IEEE Data Eng. Bull., 41:39–45, 2018. URL https://api.semanticscholar.org/CorpusID:83459546.

[19] Richard A Zann. The zebra finch: a synthesis of field and laboratory studies. Oxford University Press, 1996.

